# INSaFLU: an automated open web-based bioinformatics suite *“from-reads”* for influenza whole-genome-sequencing-based surveillance

**DOI:** 10.1101/253161

**Authors:** Vítor Borges, Miguel Pinheiro, Pedro Pechirra, Raquel Guiomar, João Paulo Gomes

## Abstract

A new era of flu surveillance has already started based on the genetic characterization and exploration of influenza virus evolution at whole-genome scale. Although this has been prioritized by national and international health authorities, the demanded technological transition to whole-genome sequencing (WGS)-based flu surveillance has been particularly delayed by the lack of bioinformatics infrastructures and/or expertise to deal with primary next-generation sequencing (NGS) data. Here, we launch INSaFLU (“INSide the FLU”), which, to the best of our knowledge, is the first influenza-specific bioinformatics free web-based suite that deals with primary data (reads) towards the automatic generation of the output data that are actually the core first-line “genetic requests” for effective and timely influenza laboratory surveillance (e.g., type and sub-type, gene and whole-genome consensus sequences, variants’ annotation, alignments and phylogenetic trees). By handling NGS data collected from any amplicon-based schema, the implemented pipeline enables any laboratory to perform advanced, multi-step software intensive analyses in a user-friendly manner without previous training in bioinformatics. INSaFLU gives access to user-restricted sample databases and projects’ management, being a transparent and highly flexible tool specifically designed to automatically update project outputs as more samples are uploaded. Data integration is thus completely cumulative and scalable, fitting the need for a continuous epidemiological surveillance during the flu epidemics. Multiple outputs are provided in nomenclature-stable and standardized formats that can be explored *in situ* or through multiple compatible downstream applications for fine-tune data analysis. This platform additionally flags samples as “putative mixed infections” if the population admixture enrolls influenza viruses with clearly distinct genetic backgrounds, and enriches the traditional “consensus-based” influenza genetic characterization with relevant data on influenza sub-population diversification through a depth analysis of intra-patient minor variants. This dual approach is expected to strengthen our ability not only to detect the emergence of antigenic and drug resistance variants, but also to decode alternative pathways of influenza evolution and to unveil intricate routes of transmission. In summary, INSaFLU supplies public health laboratories and influenza researchers with an open “one size fits all” framework, potentiating the operationalization of a harmonized multi-country WGS-based surveillance for influenza virus.

INSaFLU can be accessed through **https://insaflu.insa.pt** (see homepage view in **Figure 1**).

## Introduction

Influenza virus represents a major public health concern worldwide as it causes annual seasonal epidemics and occasional pandemics leading to high morbidity and mortality in the population (Stöhr, 2002; Iuliano et al, 2017). New viral variants emerge constantly due to the never-ending viral genetic and antigenic modification as a consequence of mutation events such as the misincorporation of nucleotides during genome replication or the exchange of genomic segments (Petrova and Russel, 2018; Westgeest KB et al, 2014). The rate of virus evolution is further shaped by the impact of the mutations on the viral fitness as well as by host immunity-related factors or ecological and environmental mechanisms, which ultimately drive the timing and frequency of the emergence of novel epidemic threats (Petrova and Russel, 2018). As such, an active molecular-based epidemiological surveillance focused on identifying patterns of viral evolution is a priority in national policies addressing influenza disease prevention, control and therapeutic measures (Petrova and Russel, 2018). To perform the genetic characterization of the virus, public health laboratories have traditionally relied on the Sanger sequencing of haemagglutinin (HA) gene, which only partially covers one of the eight negative-sense single-stranded RNA segments of the virus genome (Webster RG, 1992). Moreover, this approach almost exclusively focus the consensus sequences representing the dominant virus lineage within each infected host at a particular instant, which has limited our knowledge on intra-patient virus population diversity and transmission dynamics (Beerenwinkel et al, 2012; Dinis et al, 2016; Poon et al, 2016; Petrova and Russel, 2018). Recently, with the increased availability of next generation sequencing (NGS) technologies allowing rapid and affordable whole genome sequencing (WGS), a new era of flu surveillance has started based on genetic analysis of influenza virus at whole-genome scale (Ali et al, 2017; Revez et al, 2017; Goldstein et al, 2017;). This transition is expected to reinforce the ability of public health laboratories to: i) monitor genetic profiles of circulating influenza viruses or the emergence of pandemic influenza strains; ii) detect epitope and antiviral drug resistance mutations; iii) perform early season risk assessment; iv) strengthen vaccine effectiveness analysis; and v) optimize pre-season vaccine strain selection. In this context, there is a growing suite of influenza-specific web platforms that comprehensible allow, for instance, the annotation of phenotype-associated sequence markers, genotyping or classification of HA clades, the prediction of novel variant proteins, or even the assessment of temporal and geographical virus spread (e.g., Influenza Research Database/Fludb, Nextflu, EpiFLU/GISAID, NCBI Influenza Virus Resource, OpenFluDB) (Zhang et al, 2017; Neher and Bedford, 2015; Shu et al, 2002; Bao et al, 2008; Liechti et al, 2004). Despite their undeniable usefulness and relevance to the era of NGS-based influenza surveillance, those web-based bioinformatics tools almost exclusively rely on interrogating user-provided sequence or phylogenetic data (downstream steps). In fact, little advance has been achieved to provide public health laboratories with “influenza-specific” bioinformatics tools to deal with primary NGS data (upstream steps), which has been pointed out as the main obstacle for the demanded technological transition for flu surveillance (Ali et al, 2017). Many laboratories do not have bioinformatic capabilities and/or staff needed to timely analyze the generated NGS data (Oakeson et al, 2017; Ali et al, 2017), and, to date, NGS data has been essentially handled through in-house command-line-based pipelines or through wide multi-usage open-source (e.g., Galaxy) or commercial platforms (e.g., Geneious, CLC Genomics Workbench from QIAGEN, Bionumerics from Applied Maths or Ridom SeqSphere+ from Ridom Bioinformatics) (Shepard et al, 2016; Lee et al, 2016; Ali et al, 2017). In this context, taking advantage of the recent availability of several multiplex RT-PCR assays for whole genome amplification of influenza virus (Zhou et al, 2009; Zhou et al, 2014; McGinnis et al, 2016; Zhao et al, 2016; Ali et al, 2017; Meinel et al, 2017), we built an influenza-specific bioinformatics free web-based suite that deals with primary NGS data (reads) towards the automatic generation of the key genetic output data in a reproducible, transparent and harmonized manner that fits the diseases specificities and short-term goals for (nearly) real-time flu surveillance.

## Implementation and outputs

The bioinformatics pipeline developed and implemented in the INSaFLU web platform currently consists of 6 core steps: **1) Read quality analysis and improvement; 2) Type and subtype identification; 3) Variant detection and consensus generation; 4) Coverage analysis; 5) Alignment/phylogeny; 6) Intra-host minor variant detection (and uncovering of putative mixed infections) (Figure 2).** A summary of the INSaFLU current outputs is presented in Table 1. Documentation for each module, including software settings and current versions, is provided at the website (https://insaflu.insa.pt).

**Figure 1.**
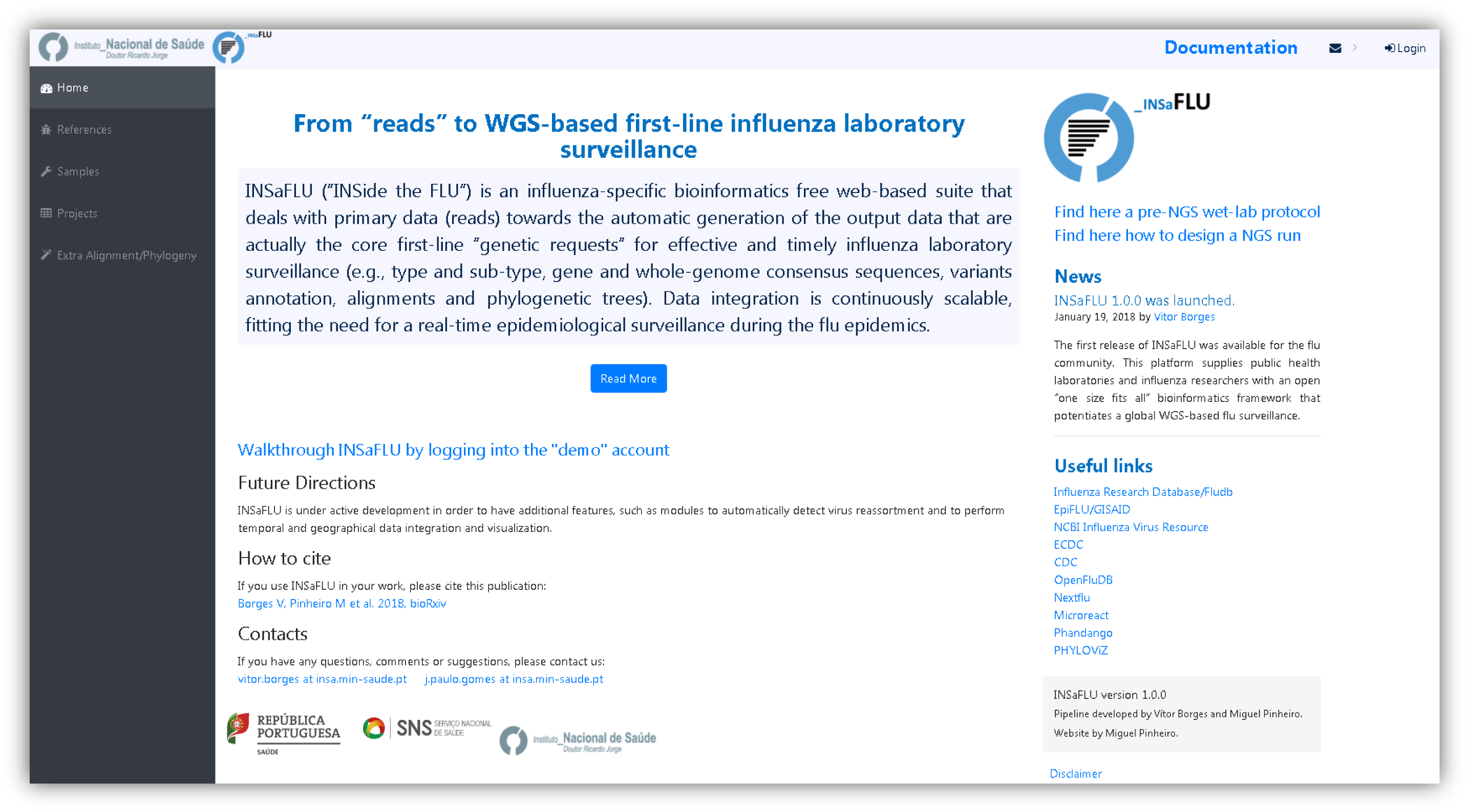
INSaFLU homepage.

**Figure 2.**
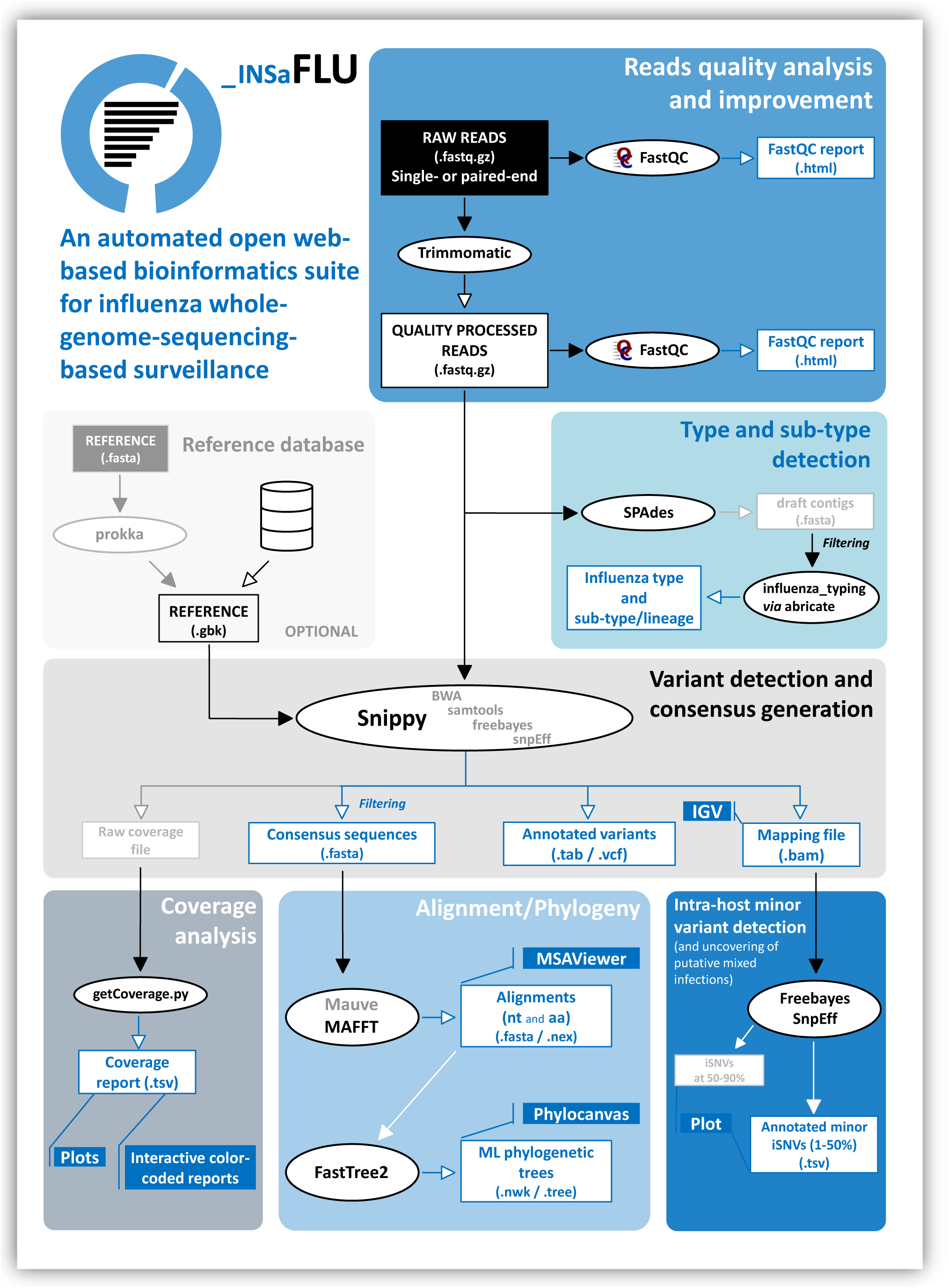
INSaFLU bioinformatics workflow.

**Table 1.**
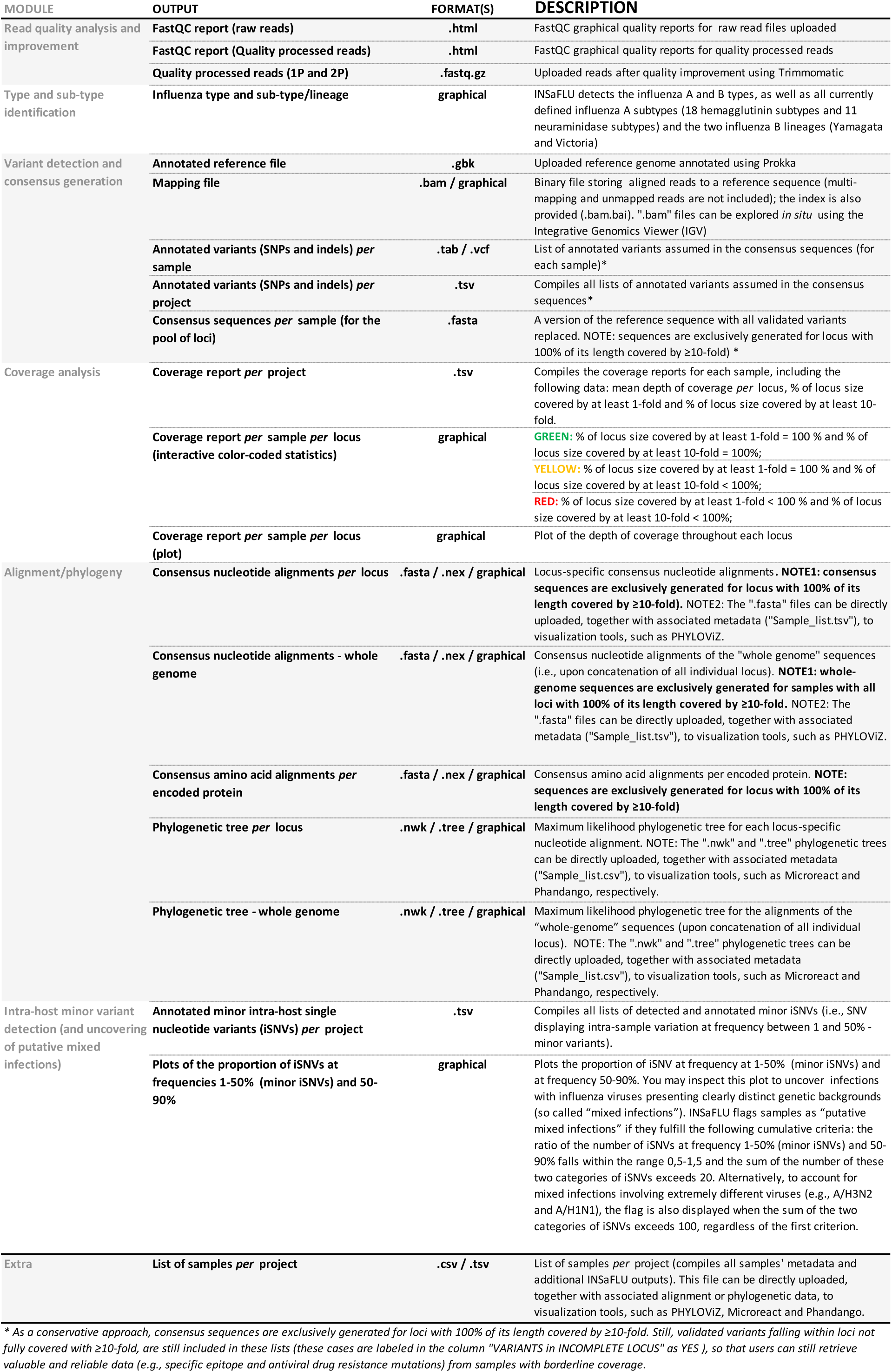
Current INSaFLU outputs.

### Read quality analysis and improvement

This module is the first step in almost all WGS bioinformatics analyses and refers to the quality control and improvement of the raw sequencing data. INSaFLU currently accepts single‐ and paired-end reads (fastq.gz format) generated through widely used NGS technologies, such as Illumina or Ion Torrent. Reads’ quality control in the INSaFLU pipeline is performed by using FastQC software (https://www.bioinformatics.babraham.ac.uk/projects/fastqc), while quality improvement is achieved through Trimmomatic (http://www.usadellab.org/cms/index.php?page=trimmomatic) (Bolger et al, 2016). This tool sequentially: i) performs a trimming sliding window by cutting reads once the average quality within a base window falls below a threshold of quality score; ii) removes very low quality bases (or N bases) from both the start and the end of each read if their quality falls below the specified minimum quality required; iii) excludes reads that fall below a specified length; and, iv) standardize the quality scores by converting them to Phred-33 scores. This first module is automatically run upon reads upload (i.e., no user intervention is needed) and provides the following outputs: i) FastQC graphical reports (“html” format) of well-established statistics of the reads quality before and after Trimmomatic analysis; and, ii) quality processed reads (“fastq.gz” format).

### Type and sub-type identification

In the second step of the pipeline (also automatically run without user involvement), a draft *de novo* assembly is performed over the quality processed reads using SPAdes (http://cab.spbu.ru/software/spades) (–*only-assembler* option) (Bankevich et al, 2012). Subsequently, the abricate tool (https://github.com/tseemann/abricate) is applied to query the draft assemblies against an *in house* database (“influenza_typing”) of a set of type‐ and sub-type/lineage-specific gene markers that allows the discrimination of the influenza A and B types, all currently defined influenza A subtypes (18 hemagglutinin subtypes and 11 neuraminidase sub-types) and the two influenza B lineages (Yamagata and Victoria). No incongruence was observed between the *in silico* determined types or HA subtypes and the result obtained by the traditional “pentaplex” real-time RT-PCR assay applied for influenza diagnosis, typing and sub-typing (Wu et al, 2013) for the tested ~250 seasonal A(H3N2) and A(H1N1pdm09) viruses. Also notable is that both or either the type and/or sub-type/lineage could be determined for viruses sequenced with very low coverage (mean depth of coverage <5-fold across the 8 amplicons), launching the perspective that this key typing data can be even retrieved from clinical samples with vestigial viruses abundance and/or generating very low PCR yield. The INSaFLU “influenza_typing” database includes: i) representative sequences of the gene encoding the matrix protein (MP or M1 gene) of influenza A and B viruses (to infer the influenza type A or B); ii) representative sequences of the HA gene of each of the 18 currently defined HA sub-types; iii) representative sequences of the neuraminidase (NA) gene of each of the 11 currently defined NA sub-types; and, iv) HA representative sequences of the influenza B lineages Yamagata and Victoria. As a proof of concept, all MP, M1, HA and NA sequences available at Influenza Virus Resource (NCBI) – Influenza Virus Database (https://www.ncbi.nlm.nih.gov/genomes/FLU/Database/nph-select.cgi?go=database), a total of 184067 sequences (database accessed in 23-25.10.2017), were screened using the INSaFLU “influenza_typing” tool. The percentage of hits correctly assigned exceeded 99,99% for NA and HA sub-typing and reached 100% for type determination. Of note, this assay detected several types/sub-types mislabeled in the NCBI database (confirmed by BLAST analyses), so these specific mis-discrepancies were not account for specificity estimation purposes. All together, upon data submission, INSaFLU automatically detects the influenza virus type and sub-type/lineage, which guides the subsequent reference-based downstream module and constitutes an optimal complement to the traditional real-time RT-PCR assays, as it discriminates any HA and NA sub-type and both influenza B lineages.

### Variant detection and consensus generation

This step of the pipeline consists of mapping the quality processed reads against user-specified reference sequences, followed by SNP/indel calling and annotation, and generation of consensus nucleotide sequences. The current reference database of INSaFLU includes reference sequences of: i) post-pandemic (2009) vaccine/reference influenza A(H1N1)pdm2009, A(H3N2) and B viruses (from both Northern and Southern hemispheres); and, ii) representative virus of multiple combinations of HA/NA subtypes (i.e., H1N1, H2N2, H5N1, H7N9, etc). All reference sequences at INSaFLU are publicly available at NCBI. The reference files, both in “.fasta” and “.gbk” (GenBank) format (annotation performed by using Prokka: https://github.com/tseemann/prokka) (Seemann et al, 2014), have been prepared to fit amplicon-based schemas capturing the whole coding sequences (CDS) of the main eight genes of influenza virus (PB2, PB1, PA, HA, NP, NA, M and NS). Nonetheless, INSaFLU is highly flexible and allows handling NGS data collected from any amplicon-based schema, provided that users fit the reference files to their amplicon design (users just have to generate and upload a multi-fasta file containing reference sequences of the individual amplicons they use with the precise size of the target sequence). Uploaded “.fasta” files are annotated using Prokka upon submission and automatically become available at the user-restricted reference database. In this module, INSaFLU takes advantage of Snippy (https://github.com/tseemann/snippy), which is a high flexible multisoftware tool for rapid read mapping (using Burrows-Wheeler Aligner – BWA; http://bio-bwa.sourceforge.net/) (Li and Durbin, 2009), SNP‐ and indel calling (using samtools: http://samtools.sourceforge.net/; and freebayes: https://github.com/ekg/freebayes) (Li et al, 2009; Garrison and Marth, 2012), variant annotation (using SnpEff: http://snpeff.sourceforge.net/) (Cingolani et al, 2012) and consensus generation (using vcftools: http://vcftools.sourceforge.net/) (Danecek et al, 2011). We selected the following criteria for reads mapping and validating SNPs /indels to be annotated, listed and assumed in the consensus sequences: i) a minimum mapping quality of ≥20; ii) a minimum number of 10 quality processed reads covering the variant position; iii) a minimum proportion of 51% of quality processed reads at the variant position differing from the reference. As a conservative approach, for each virus, consensus sequences are exclusively generated for loci with 100% of its length covered by ≥10-fold (see below the “Coverage analysis” module for more details), thus avoiding the generation of incomplete sequences that would shrink the nucleotide region available for genetic diversity analyses. Nonetheless, variants that fulfill the above described criteria, but fall within loci not fully covered with ≥10-fold, are still included in the list of all variants per sample/project (a specific flag is provided for these cases), so that users can still retrieve valuable and reliable data (e.g., specific epitope and antiviral drug resistance mutations) from samples with borderline coverage. Users can explore all output mapping files (".bam" format) to view and inspect all reads and variants using the easy-to-use visualization tool Integrative Genomics Viewer (http://software.broadinstitute.org/software/igv/) (Robinson et al, 2011) available at INSaFLU. These output files are also used in INSaFLU pipeline to more complex downstream analyses (see below the module “Intra-host minor variant analyses”). For each run (see INSaFLU usage section), users must choose the reference sequences (in general, the vaccine reference sequences of the season under surveillance) and the pool of samples to be compared (viruses sharing the same type/subtype as the reference selected, as inferred in the previous module). The option to map reads against same type and sub-type reference sequences of the vaccine reference strains not only potentiates the mapping quality, but also has the clear advantage of providing the user with a list of amino acid replacements properly coded to be reported for surveillance. In fact, the amino acid substitutions (including key markers of specific clades/genetic groups) that are reported by National Reference Laboratories to supranational health authorities (e.g., reports to ECDC/WHO via TESSy) are coded against the sequence profile of vaccine strains. In summary, this INSaFLU module provides the key data that are actually the core first-line "genetic requests" for effective and timely monitoring of influenza virus evolution on behalf of seasonal influenza laboratory surveillance, i.e. the list of variants (assumed in consensus sequences) and their effect at protein level and also consensus sequences. The latter constitutes the whole basis for the downstream phylogenetic inferences driving the continuous tracking of influenza temporal/geographical spread.

### Coverage analysis

A key standard parameter to take into account when performing NGS is the mean depth of coverage, defined as the mean number of times each base shows up in individual reads (also known as vertical coverage). When handling small amplicon-based NGS data for virus variant detection and consensus generation, it is mandatory to finely inspect the fluctuation of the depth of coverage throughout each amplicon region (Beerenwinkel et al, 2012). Such inspection of the so-called “horizontal coverage” may not only be highly informative about sequencing-derived artifacts (the coverage plot should typically follow an invert U shape *per* amplicon), but also provides important clues about the degree of relatedness between the genetic background of the “query” virus and the reference sequence chose for mapping. For instance, obtaining sufficient mean depth of coverage for a given amplicon for which its complete length was not covered at 100% may be indicative of miss-mapping due to a high genetic distance between the reference sequence for that locus and the virus under sequencing. These phenomena are typically expected for cases of antigenic shift (reassortment between viral segments from different strains) or intra-segment homologous recombination, or even, for instance, for cases of “mis-subtyping” or “mis-choice” of the reference sequences (e.g. erroneous mapping of A/H1N1pdm09 viruses against a vaccine A/H3N2 reference). In this context, we developed the *getCoverage.py* script (available at https://github.com/monsanto-pinheiro/getCoverage), so that INSaFLU automatically provides the user with a deep analysis of the coverage both *per* sample (graphical outputs) and as batch *per* project (“.tsv” format), by yielding the following data: mean depth of coverage *per* locus, % of locus size covered by at least 1-fold and % of locus size covered by at least 10-fold. The latter statistics was chosen both to fit the minimum depth of coverage for variant calling and to guide the consensus generation (as described above), i.e., the consensus sequences are exclusively provided for amplicons fulfilling the criteria of having 100% of their size covered by at least 10-fold. In addition, INSaFLU interactively yields intuitive color-coded outputs of the coverage statistics as well as depth of coverage plots for each locus *per* sample (see example in Figure 3), enabling users to fine-tune this important parameter towards the uncovering of eventual atypical but highly relevant genetic events, such as reassortment/homologous recombination events.

**Figure 3.**
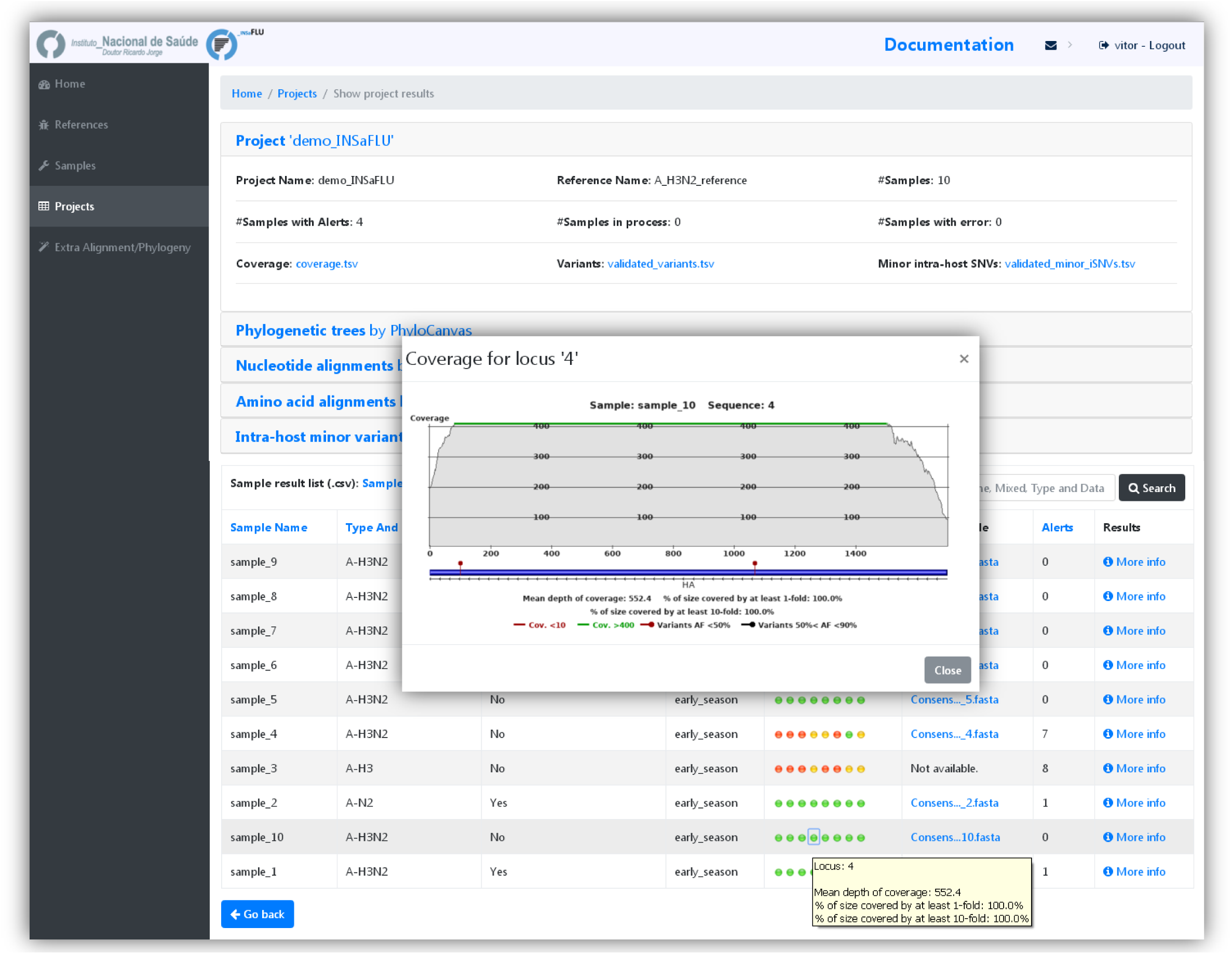
Coverage inspection.

### Alignment/phylogeny

On behalf of the continuous tracking of influenza virus evolution during epidemics, it is mandatory to apply bioinformatics tools that facilitate a deep exploration and analysis of the genetic diversity of the circulating virus. To the best of our knowledge, there is no upstream influenza-specific bioinformatics web tool to deal with primary NGS data towards the generation of harmonized sequence or phylogenetic data that can be directly applied for such fine-tune downstream analysis and visualization platforms. INSaFLU was specifically designed to overcome this caveat in order to obviate the operationalization of a harmonized supranational WGS-based surveillance of influenza virus (ECDC, 2016; Ali et al, 2017). In this module, filtered consensus nucleotide sequences are used as input to progressiveMAUVE (http://darlinglab.org/mauve/mauve.html) (Darling et al, 2010) and MAFFT (https://mafft.cbrc.jp/alignment/software/) (Katoh et al, 2002) for draft and subsequent refined sequence alignment, respectively. INSaFLU provides refined nucleotide sequence alignments (FASTA and NEXUS formats) both at locus level, i.e. for each one of amplicon targets (which are, in general, influenza CDSs), and at “whole-genome” scale (after concatenation of all amplicon targets). Amino acid alignments for annotated proteins are also built using MAFFT (Katoh et al, 2002). Phylogenetic trees (in standard ".nwk" and ".tree" formats) are inferred for each alignment by maximum likelihood under the General Time-Reversible (GTR) model (1000 bootstraps) using double-precision mode of FastTree2 (
https://www.microbesonline.org/fasttree/) (Prince et al, 2010). In order to fulfill the demands of the cumulative data acquisition underlying laboratory surveillance throughout each flu season, for each INSaFLU project, alignments and phylogenetic trees are automatically re-build and updated as more samples are added, making data integration completely flexible and scalable (see "Usage" section). Alignments and phylogenetic trees can be either downloaded for external exploration or explored *in situ* at INSaFLU website using MSAViewer (http://msa.biojs.net/) (Yachdav et al, 2016) and PhyloCanvas (http://phylocanvas.org/), respectively. Moreover, INSaFLU will soon enable running this module in an independent manner over a set of user-selected sequences, so users are also encouraged to upload (to a user-restricted reference database) gene or whole-genome sequences of representative virus of specific genetic (sub)groups/clades/lineages, so that phylogenetic diversity of circulating viruses can be better evaluated and integrated in the frame of guidelines defined by supranational health authorities (e.g., ECDC) for each season.

In summary, INSaFLU dynamically builds ready-to-explore scalable gene‐ and genome-based alignments (see example in Figure 4) and phylogenetic trees (see example in Figure 5) in standardized nomenclatures and formats that fully compatible with multiple downstream applications. These include not only other web-based "surveillance-oriented" platforms for influenza genotyping, phenotypic prediction or phylogeographical/patient data integration (such as, PHYLOViZ, Phandango and Microreact) (Ribeiro-Gonçalves et al, 2016; Hadfield et al, 2017; Argimón et al, 2016), but also several computationally intensive bioinformatics algorithms commonly applied for fine-tune research of influenza evolutionary dynamics, such as inference of signatures of selection or refined phylogenetics (e.g. the widely used MEGA, DnaSP, BEAST and RAxML).

**Figure 4.**
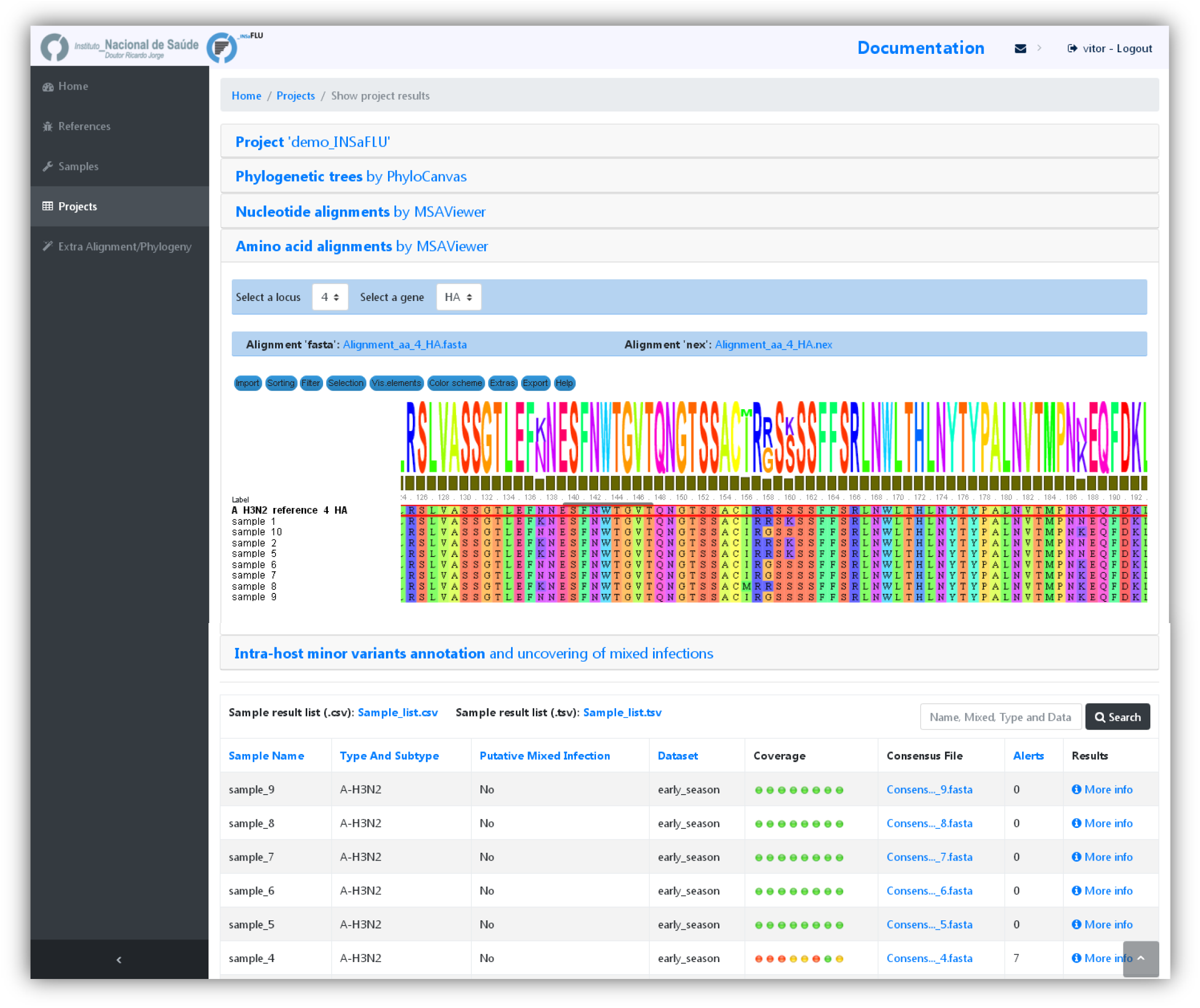
Alignment visualization.

**Figure 5.**
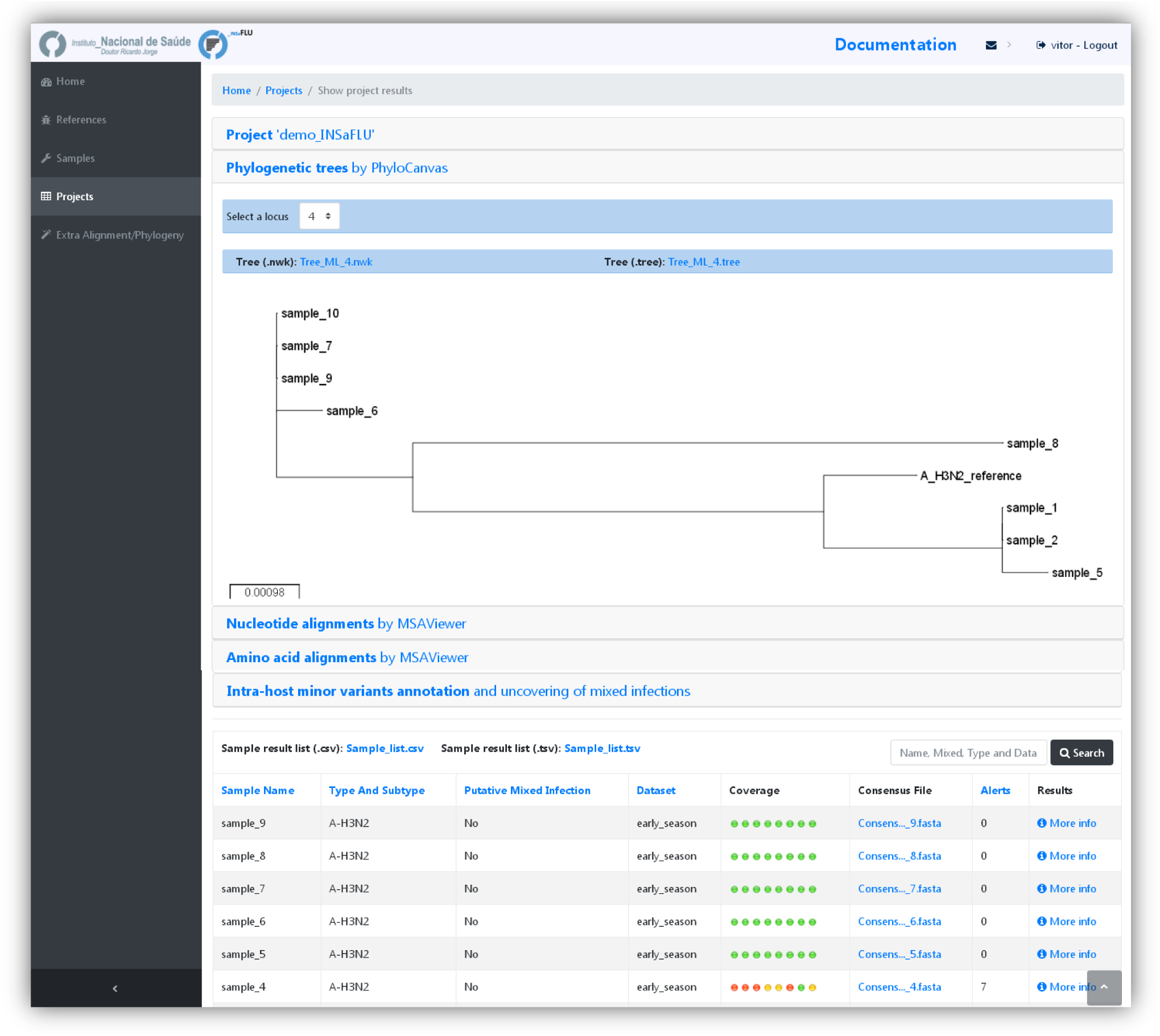
Phylogenetic trees visualization.

### Intra-host minor variant detection (and uncovering of putative mixed infections)

As a result of the huge advances in in-depth NGS technologies and associated bioinformatics, a currently growing research interest on influenza field relies on investigating the virus evolutionary dynamics within the human host during infection. In this context, INSaFLU additionally provides the user the possibility to get insight on the influenza intra-patient sub-population dynamics through the scrutiny of minor intra-host single nucleotide variants (iSNVs), i.e., SNV displaying intra-sample frequency below 50%. This is achieved by applying *freebayes* software (https://github.com/ekg/freebayes) (Garrison and Marth, 2012) over mapping files (“.bam” format) with the following criteria: i) excludes read alignments from analysis if they have a mapping quality of less than 20; ii) excludes alleles from iSNV analysis if their supporting base quality is less than 20; iii) requires a minimum of 100-fold depth of coverage to process a site for iSNV analysis; and, iv) requires at least 10 reads supporting an alternate allele within a single individual to evaluate the iSNV frequency. Once fulfilling the above previous criteria, no less than 1% of intra-host frequency of the alternate allele is reported. As such, in a dynamic manner, distinct minimum iSNV frequency cut-offs are assumed depending on the depth of coverage reached at each site, i.e., the identification of iSNV sites at frequencies of 10%, 2% and 1% is only allowed if the depth of coverage at a particular site exceeds 100-fold, 500-fold and 1000-fold, respectively. For each INSaFLU project, results are compiled in a table (“.tsv” format) listing all iSNVs (detected for all project’ samples) at frequencies between 1% and 50%. As above, variant annotation (using SnpEff: http://snpeff.sourceforge.net/) (Cingolani et al, 2012) is also provided. Of note, variants at a frequency above 50%, which correspond to variants included in the consensus sequences, are filtered out from this table since they are systematically listed and annotated downstream in the pipeline (see module “Variant detection and consensus generation”). The table can easily be scrutinized to find sites displaying inter-patient redundancy (i.e., iSNV sites found in more than one individual). These may for instance constitute the ultimate genetic clues for disclosing influenza transmission links (Poon et al, 2016) or the emergence of antiviral resistance (Flaherty et al, 2012; Trebbien et al, 2017). Similarly to what is outlined in the previous module, this table is automatically re-build and cumulatively updated as more samples are added to each INSaFLU project. In order to additionally enable the detection of infections with influenza viruses presenting clearly distinct genetic backgrounds (so called “mixed infections”), INSaFLU additionally plots the proportion of iSNV at frequency 1-50% (minor iSNVs) and 50-90% detected for each sample (the positional mapping of iSNVs from these two categories within each amplicon can also be explored in the “coverage plots”; see above). A cumulative high proportion of iSNVs at both frequency’ ranges is mostly likely to represent a mixed infection, in a sense that the natural intra-patient influenza diversification (that NGS is capable of detecting) is expected to be very low (no more than a few tenths of variants, most of them at frequency <10%) (Debbink et al, 2017; Poon et al, 2016; Dinis et al, 2016). INSaFLU flags samples as “putative mixed infections” if they fulfill the following cumulative criteria: the ratio of the number of iSNVs at frequency 1-50% (minor iSNVs) and 50-90% and falls within the range 0,5-1,5 and the sum of the number of these two categories of iSNVs exceeds 20. Alternatively, to account for mixed infections involving extremely different viruses (e.g., A/H3N2 and A/H1N1), the flag is also displayed when the sum of the two categories of iSNVs exceeds 100, regardless of the first criterion. These numerical indicators were inferred upon multiple testing, including the independent NGS run of sample replicates constituting “true” mixed infections (see example in Figure 6).

**Figure 6.**
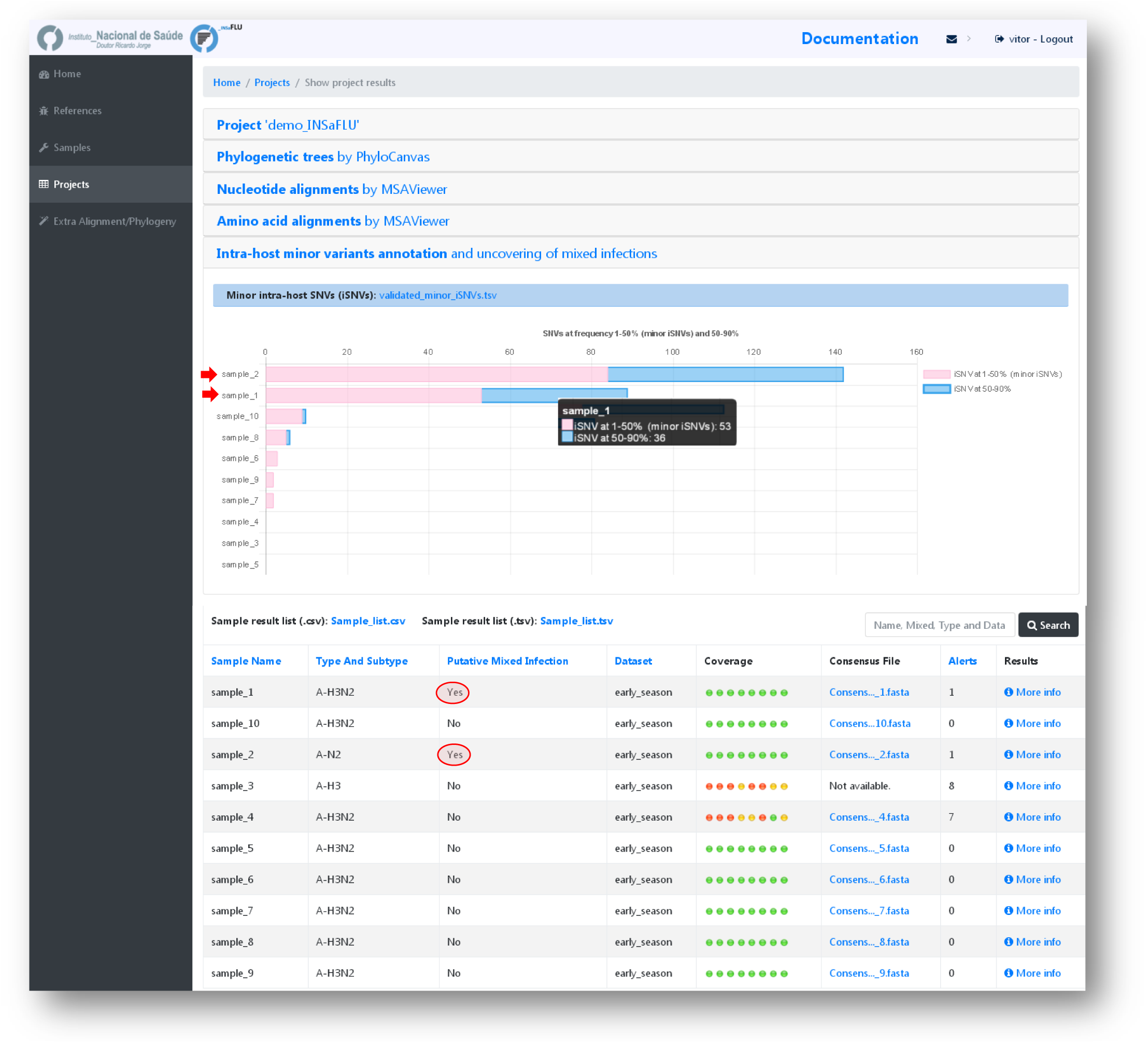
Intra-host minor variant detection (and uncovering of putative mixed infections).

In summary, through this module, INSaFLU supplies public health laboratories and influenza researchers with relevant data on influenza sub-population diversification within humans that can be systematically integrated in parallel with the “classical” data on “consensus-based” inter-patient virus genetic diversity. Taking into account the recent findings on this subject (Flaherty et al, 2012; Poon et al, 2016; Trebbien et al, 2016; Sobel Leonard et al, 2016; Dinis et al, 2016; Debbink et al, 2017; McCrone et al, 2017), it is expected that this dual approach will strengthen not only our ability to detect the emergence of antigenic and drug resistance variants, but also to decode alternative pathways of influenza evolution and to unveil intricate routes of transmission.

In summary, through this module, INSaFLU supplies public health laboratories and influenza researchers with relevant data on influenza sub-population diversification within humans that can be systematically integrated in parallel with the “classical” data on “consensus-based” inter-patient virus genetic diversity. Taking into account the recent findings on this subject (Flaherty et al, 2012; Poon et al, 2016; Trebbien et al, 2016; Sobel Leonard et al, 2016; Dinis et al, 2016; Debbink et al, 2017; McCrone et al, 2017), it is expected that this dual approach will strengthen not only our ability to detect the emergence of antigenic and drug resistance variants, but also to decode alternative pathways of influenza evolution and to unveil intricate routes of transmission.

## Pre-NGS design and full pipeline testing

The INSaFLU pipeline has been mainly tested with NGS data (192 samples from the pilot season 2016/2017) generated in an Illumina MiSeq apparatus after influenza whole genome amplification with a modified wet-lab protocol based on a previously reported RT-PCR assay (Zhou et al, 2009, for Influenza A; and Zhou et al, 2014, for Influenza B; Zhou and Wentworth DE, 2012). The adapted pre-NGS protocols, both for influenza A and B viruses, are provided in the INSaFLU’s documentation and can be straightforwardly used for the routine generation of amplicon template for WGS of influenza viruses (irrespective of virus sub-type / lineage). Library preparation was conducted following the Nextera XT DNA Library Prep Reference Guide and WGS runs (96 samples per run) were carried out using MiSeq Illumina flow cells to obtain 2 × 150 paired-end reads (300 cycles). Based on our experience with the described experimental design, success (i.e., 100% of the length of the 8 influenza CDS covered by ≥ 10-fold) is largely potentiated if WGS runs are designed to yield >150000 (2 × 75000) reads *per* sample. In fact, above this cut-off, a success of 92% was achieved when comparing with less than 70% obtained for samples with <150000 dedicated reads. As a prudent approach, users should design NGS runs to go further this cut-off (e.g., 300000 reads per sample) in order to better account for issues arising from both the PCR (e.g., fluctuations in the percentage of influenza-specific amplicons across samples and unbalanced relative proportions of the in-sample amplicons) and the NGS run (e.g., low yield and unbalanced demultiplexing of the reads across the samples).

## Usage

INSaFLU is a free of charge platform located at the website of the Portuguese National Institute of Health, Instituto Nacional de Saúde (INSA) Doutor Ricardo Jorge. It can be openly used upon account creation. This allows data storage/update in a continuous manner, thus facilitating continuous epidemiological surveillance. INSaFLU gives access to private sample and reference databases and projects’ management. All data is user-restricted, so it will not be viewable by other users.

All that is really needed to use INSaFLU is a computer with connection to the Internet. A tutorial providing a complete usage example of data upload, project launching and management, as well as of how to visualize/download graphical and sequence/phylogenetic output data is provided at INSaFLU’s DOCUMENTATION (http://insaflu.readthedocs.io/). Users can also walkthrough INSaFLU by logging into a “**demo**” account (https://insaflu.insa.pt/accounts/login/). The web platform architecture is quite intuitive and enrolls the following main tabs: **Samples, References and Projects.**

<u>**Samples.**</u> This menu displays all information for all samples loaded by the user. Required sample-associated data to be uploaded at INSaFLU include:

- **NGS data**: single‐ or paired-end reads (fastq.gz format) obtained through NGS technologies, such as Illumina or Ion Torrent (reads can be submitted individually or as a batch);
- **Sample metadata**: a table file can be uploaded for a batch of samples (preferable option) or the sample’s information can be inserted individually at the INSaFLU platform. In order to link the sample data to the uploaded read files, the table file [in comma-separated value (csv) or tab-separated value (tsv)] should contain the columns “sample name”, “fastq1”, “fastq2” (mandatory columns to fulfill; "fastq2" is exceptionally not fulfilled for single-end data) as well these additional variables (that may not be fulfilled), which commonly constitute the typical metadata collected during seasonal influenza surveillance: "data set", "vaccine status", "week", "onset date", "collection date", "lab reception date", "latitude", "longitude". However, users may include any other columns with metadata variables to be associated with samples. An example table file is provided at the website. The option to upload tables enriched with multiple metadata variables has the clear advantage of allowing their subsequent direct upload (along to the standardized and multi-format outputs of INSaFLU: alignments/trees) to downstream platforms for phylogenetic data visualization and/or phylogeographical analysis, such as PHYLOViZ (http://www.phyloviz.net) (Ribeiro-Gonçalves, 2016), which accepts sample metadata (tab-separated format) plus alignments (FASTA format), Phandango (https://jameshadfield.github.io/phandango/#/) (Hadfield et al, 2017), which runs sample metadata (csv-separated format) and a phylogenetic tree (“.tree” format) or Microreact (https://microreact.org/) (Argimón et al, 2016), which takes sample metadata (in csv-separated format) plus a phylogenetic tree (“.nwk” format).

Upon submission, INSaFLU automatically updates samples information with read’s quality and typing data.

<u>**References.**</u> This menu displays all information for all reference sequences available at user's confidential account. INSaFLU provides a default reference database including publicly (NCBI) available sequences from several post-pandemic (2009) vaccine/reference virus and representative virus of multiple combinations of HA/NA subtypes. The database includes both locus and whole-genome sequences (FASTA and GenBank formats) that are ready to be used for reference-based mapping (Project type “Whole workflow”) or for alignment/phylogeny analyses (Project type “Extra Alignment/Phylogeny”) (see next section). Nonetheless, users are allowed to upload additional reference files to a user-restricted reference database (uploaded “.fasta” files are automatically annotated upon submission).

<u>**Projects.**</u> Two types of projects can be launched at INSaFLU: whole workflow *Projects* (to run the complete pipeline) or *Extra Alignment/Phylogeny* projects (to perform additional genetic diversity analyses).

- **Whole workflow *Project*.** Upon the project creation, users must select: i) a reference file from the reference database that fit their amplicon design (i.e., a multi-fasta file containing reference sequences of the individual amplicons they use with the precise size of the target sequence); and ii) the batch of samples to be included in the project. Since the projects are scalable, users are encouraged to create “umbrella” projects, such as projects enrolling the mapping of all same subtype virus against the vaccine reference virus for a given flu season. Outputs of the project are organized by dynamic “expand-and-collapse” panels that allow a user-friendly visualization/download of all graphical and sequence output data.
- **Extra Alignment/Phylogeny** projects (<u>available soon</u>). The rationale behind the usage and management of these projects is the same as the one described for “whole workflow” projects. Here, users must select sequences (same-locus sequences or whole-genome sequences) from the reference database and/or from an INSaFLU sequence repository that includes all validated consensus sequences generated by the several user-restricted “whole workflow” projects. This module provides a huge flexibility in a sense that allows that the genetic diversity of circulating viruses can be better evaluated and integrated according with the users’ specific needs. For instance, users may perform extra gene‐ or whole-genome-scale alignment/phylogenetic analyses enrolling: i) not only the circulating viruses, but also representative virus of specific genetic (sub)groups/clades/lineages as defined by supranational health authorities guidelines (e.g., ECDC) for each season; ii) viruses from different seasons; iii) particular subsets of samples from a given project (e.g., subset of viruses from a specific genetic clade or subsets excluding samples flagged as “putative mixed infection”), etc.

Finally, it is worth noting that INSaFLU relies on a multi-software bioinformatics pipeline that will be under continuous development and improvement not only to enrich it with new features, but also to continuously shape the pipeline to the best bioinformatics advances in the field. The current software settings were chosen upon intensive testing and are detailed for consultation at website (DOCUMENTATION menu). Although pipeline changes will be systematically kept available for easy historical consultation, software versions are provided for every INSaFLU run.

## Conclusion

Most Reference Laboratories for influenza surveillance are still not capacitating to handle the NGS data, which has delayed the demanded technological transition to a WGS-based flu surveillance (ECDC, 2016; Ali et al, 2017). In this regard, we developed INSaFLU, which, to the best of our knowledge, is the first influenza-specific bioinformatics open web-based suite that deals with primary NGS data (reads) towards the automatic generation of the output data that are actually needed for the first-line influenza surveillance (type and sub-type, gene and whole-genome sequences, alignments and phylogenetic trees). The main advantages offered by INSaFLU are:

i. it allows handling NGS data collected from any amplicon-based schema;
ii. it enables laboratories to perform advanced, multi-step software intensive analyses in a user-friendly manner without previous training in bioinformatics;
iii. it is free of charge and can be used upon account creation giving access to user-restricted sample and reference databases and projects’ management;
iv. it is located at the website of a National Institute of Health, which ensures confidentiality and ethics;
v. it is a flexible tool specifically designed to integrate output data in a cumulative manner, thus fitting the analytical dynamics underlying a continuous epidemiological surveillance during the flu epidemics;
vi. outputs are provided in nomenclature-stable and standardized format and can be explored *in situ* or through multiple compatible downstream applications for fine-tune data analysis.

In summary, INSaFLU provides an open “one size fits all” framework that guarantees that the ultimate technology for flu surveillance can be easily accessed by any laboratory around the world with a common computer with access to Internet. It is launched to potentiate the operationalization of a harmonized multi-country WGS-based surveillance for Influenza virus. It will certainly strengthened the detection of genetic changes in circulating influenza viruses, the detection of potential pandemic influenza strains, the early season risk assessment and vaccine effectiveness analysis, the detection of genetic markers associated with antiviral resistance, and pre-season vaccine strain selection. Ultimately, INSaFLU has the potential to obviate collaborative initiatives among cross-sectorial stake-holders enrolled in the flu surveillance, with benefits for public health.

## Future Directions

INSaFLU is under active development in order to have additional features, such as modules to automatically detect virus reassortment, and to perform temporal and geographical data integration and visualization.

## Acknowledgements

We thank to Joana Mendonça and Luís Vieira from the Innovation and Technology Unit of the Department of Human Genetics of the Portuguese National Institute of Health for their support on NGS runs.

## Funding

*Conflict of interest: none declared*

